# Investigating individual variability in microstructural-functional coupling in the human cortex

**DOI:** 10.1101/2023.05.29.542730

**Authors:** Raihaan Patel, Alyssa Dai, Sofie L. Valk, Gabriel Desrosiers-Grégoire, Gabriel A. Devenyi, M. Mallar Chakravarty

**Affiliations:** Cerebral Imaging Centre, Douglas Mental Health University Institute, Verdun, Canada; Department of Biological and Biomedical Engineering, McGill University, Montreal, Canada; Integrated Program in Neuroscience, McGill University, Montreal, Quebec, Canada; Max Planck Institute for Human Cognitive and Brain Sciences, Leipzig, Germany; INM-7, Forschungszentrum Jülich, Jülich, Germany; Department of Psychiatry, McGill University, Montreal, Canada

## Abstract

Understanding the relationship between the structural and functional architecture of the human brain remains a key question in neuroscience. In this regard variation in cortical myelin may provide key insights into the functional organization. Previous findings have demonstrated that regions sharing myeloarchitectonic features are also likely to be structurally and functionally connected. However, this association is not uniform for all regions. For example, the strength of the association, or ‘coupling’, between microstructure and function is regionally heterogeneous, with strong coupling in primary cortices but weaker coupling in higher order transmodal cortices. However, the bases of these observations have been typically made at the group level, leaving much to be understood regarding the individual-level behavioural relevance of microstructural-functional coupling variability. To examine this critical question, we apply a multivariate framework to a combination of high-resolution structural, diffusion, and functional magnetic resonance imaging (MRI) data in a sample of healthy young adults. We identify four distinct patterns of coupling variation that vary across individuals. Remarkably, we find that while microstructural-functional coupling is consistently strong in primary cortices, significant variation in transmodal cortices exists. Importantly, we identified coupling variability maps and their association with behaviour that demonstrate the existence of latent dimensions of variability related to inter-individual performance on cognitive tasks. These findings suggest that the existence of behaviourally relevant coupling variation is a key principle for brain organization.

## 1. Introduction

The relationship between brain structure and function is a central question in neuroscience research. In this regard, cyto- and myelo-architectonic principles have been used to describe local architecture properties that can then be used to parcellate the brain (Amunts & Zilles, 2015; Lerch et al., 2017; Nieuwenhuys & Broere, 2020). Importantly, the similarity between brain parcels have been used to define brain connectivity at the single subject and population level (Barbas, 2015; Eickhoff et al., 2007; García-Cabezas et al., 2019; Hilgetag et al., 2019; Pandya et al., 2015; Valk et al., 2020; Yeo et al., 2011; Zikopoulos et al., 2018). Multiple lines of research suggest that the cyto- and myelo-architectonic similarity of two brain regions may relate to axonal connectivity, even if regions are separated by large distances (Barbas, 2015; García-Cabezas et al., 2019). These findings are confirmed in studies of model organisms where cellular and molecular examinations are more attainable. For example, in the macaque (Barbas & Rempel-Clower, 1997; Beul et al., 2017) and mouse (Goulas et al., 2017), post-mortem histology and tract tracing techniques have been used to demonstrate that cytoarchitectonic similarity is indicative of inter-areal anatomical connections. However, studying the relationship between various domains of brain architecture and their relationship to behaviour remains an active and important area of investigation.

To this end, recent and continuing advances in high quality magnetic resonance imaging (MRI) datasets have enabled study of these principles *in vivo* in the human brain. Using a number of MRI derived metrics it is now possible to probe brain microstructure sensitive to to neuronal density, fiber organization, and myelin concentration (Edwards et al., 2018; Lerch et al., 2017; Olafson et al., 2021; Paus, 2018; Tardif et al., 2016). The integration of function, as defined using resting state functional MRI (rsfMRI) (Biswal et al. 1995; Smith et al. 2009; Smith et al. 2013; Fox and Greicius 2010; Greicius et al. 2003) and microstructure provides the potential to examine the extent to which local microstructural architecture relates to functional brain organization *in vivo*. In this context, previous work demonstrated that intracortical myelin (assessed via quantitative T1) is positively associated with functional connectivity (as measured using rsfMRI) (Huntenburg et al., 2017).

Coupling between microstructure and function demonstrates marked topographic dependencies. When assessed in the context of a functional gradient (Margulies et al., 2016) microstructure-function coupling strength is increased in unimodal regions such as motor and visual areas, but this correspondence weakens moving towards transmodal association areas (Huntenburg et al., 2017; Paquola et al., 2019; Sydnor et al., 2021). Therefore, study of the spatial variations of microstructure-function coupling may have overarching implications for brain organization.

An open question in the context of structure-function coupling and decoupling, particularly in transmodal cortices, is whether it is related to inter-individual differences (Suárez et al., 2020). The impact of this variation cannot easily be elucidated from previous studies as the methodologies used typically describe the group-level relationship of structure-function coupling, thereby obscuring potentially relevant inter-individual variation in behavior (Mueller et al., 2013). In this work, we aim to investigate the existence and behavioural relevance of inter-individual variability in microstructure-function coupling across the human cortex using high-resolution structural and functional MRI data from the Human Connectome Young Adult

Dataset (Van Essen et al., 2013). We measure microstructure via the recently developed microstructural profile covariance (MPC) approach which samples multiple cortical depths at each point along the cortical surface in myelin sensitive MRI data (Paquola et al., 2019). We examine the relationship of these microstructural profiles to functional connectivity by generating modes of variation using non-negative matrix factorization (NMF) (D. D. Lee & Seung, 1999; D. Lee & Seung, 2001). NMF has been used previously by our group (Patel et al., 2022; Patel, Steele, Chen, Patel, Devenyi, Germann, Tardif, & Mallar Chakravarty, 2020; Robert et al., 2021) and others (Nassar et al., 2018; Sotiras et al., 2015) to localize inter-individual differences. Our findings identify novel models of microstructural-functional coupling that may underlie basic brain organization and behaviour.

## 2. Methods

### 2.1 Overview

We used structural and functional MRI data from a sample of 384 unrelated individuals from the Human Connectome Project (HCP) Young Adult Dataset (Van Essen et al., 2013) (**2.2** Data) to avoid potential influences of family structure and genetic bias. Microstructural profile covariance (MPC) was computed between each pair of 400 brain regions, for each subject (**2.2** Microstructural Profile Covariance Computation). Similarly, resting state functional connectivity was computed between each pair of 400 brain regions, for each subject (**2.3** Resting State Functional Connectivity). We compute a coupling coefficient for each of the 400 brain regions representing the correlation of a region’s functional connectivity to all other regions with its microstructural similarity to all other regions (**2.4** Computing structure-function coupling, **Figure 1A**). We decompose the coupling data for all subjects into linear components using NMF (**2.6** Identifying coupling components with NMF, **Figure 1B**), and assess behavioural relevance of coupling variation using partial least squares (PLS) (**2.7** Identifying the cognitive determinants of Structure-Function Coupling, **Figure 1C**). Code for this work is available at https://github.com/raihaan/hcp-micro-func.

**Figure 1.**
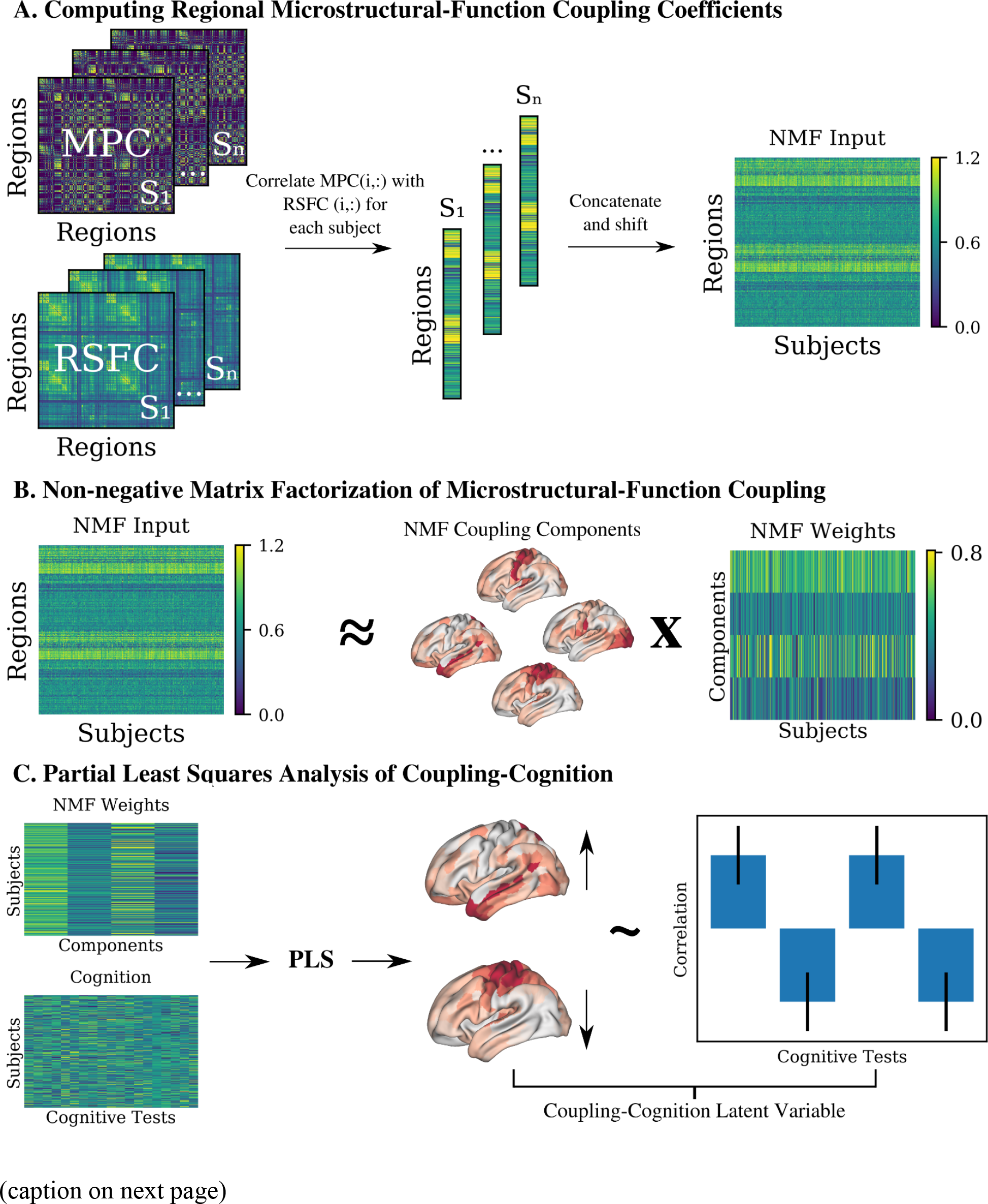
Overview of the methodological approach employed. (A) Node level microstructural-functional coupling coefficients are computed by correlating each row of MPC matrix with the corresponding row of the RSFC matrix on an individual subject basis. Coupling coefficients are stacked in columnar fashion to form the NMF input matrix. (B) NMF decomposes the input coupling data into a set of additive spatial components and subject weightings. (C) PLS analysis identifies latent variables which maximize covariance between NMF weightings and cognitive performance, thus identifying the multivariate relationship between coupling variability and variability in cognitive performance.

### 2.2 Microstructural Profile Covariance Computation

MPC was computed via comparison of cortical depth profiles of T1w/T2w ratio data, as in (Paquola et al., 2019; Valk et al., 2022) (see Supplementary Materials, Section 1.2 for details). The ratio of T1w/T2w has been shown to recapitulate known patterns of cortical myelin and microstructure (Ganzetti et al., 2014; Glasser & Van Essen, 2011; Paquola et al., 2019), as well as to track gradients of laminar differentiation (García-Cabezas et al., 2020). The MPC was estimated using Freesurfer version 5.3-HCP generated pial and grey-white matter cortical surfaces. The MPC is estimated from the intensity profile generated per vertex across 12 surfaces generated across the cortex using the equivolume approach (Waehnert et al., 2014). For a given vertex on the cortical surface, this presents the opportunity to sample at multiple cortical depths as opposed to collapsing the full columnar variation onto a single point. We sampled T1w/T2w at each of the 12 surface-specific vertices to obtain a microstructural profile of each vertex. We applied the Schaefer 400 parcellation (https://github.com/ThomasYeoLab/CBIG/tree/master/stable_projects/brain_parcellation/Schaefer2018_LocalGlobal) (Schaefer et al., 2018) in order to obtain the average microstructural profile in each of 400 cortical regions. The MPC between each pair of regions *i* and *j* is then computed as defined previously (Paquola et al., 2019):

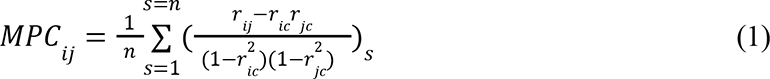

where *r_ic_* is the correlation of the microstructural profile of region *i* with the average microstructural profile across the whole brain, *r_jc_* is the correlation of the microstructural profile of region *j* with the average microstructural profile across the whole brain, *s* is a given participant and *n* is the number of subjects. Here we compute MPC on a per subject basis, such that *n*=1 and the above is calculated for each individual subject separately. The result of this process is a matrix MPC with dimensions 400 × 400 being computed for each individual subject.

### 2.3 Resting State Functional Connectivity

We computed resting state functional connectivity (RSFC) matrices for each subject, derived from the HCP’s minimally preprocessed fMRI data. This includes corrections for gradient non-linearities and motion, with reverse phase encode acquisitions used to compute and correct for geometric distortions (Glasser et al., 2013). Corrected data was then warped to MNI space where bias field correction, brain extraction, and a high pass filter was applied to correct for scanner drift (Glasser et al., 2013). ICA-FIX, an automated classifier to detect and remove non-neuronal noise sources, was then applied (Salimi-Khorshidi et al., 2014). Finally, rsfMRI data is sampled onto the cortical mid-surface of MSMALL aligned cortical surfaces of each subject (Robinson et al., 2014). We did not perform global signal regression, based on recent evidence suggesting that the global signal may contain biologically relevant information (Hyder & Rothman, 2010; Li et al., 2019; Schölvinck et al., 2010; Wong et al., 2013) as well as a current lack of consensus (Bijsterbosch et al., 2020; Liu et al., 2017; Murphy & Fox, 2017; Saad et al., 2012). Each of two 15 minute resting state sessions were variance normalized and then concatenated to form a single 30 minute time series for each individual. We next applied the Schaeffer parcellation to obtain the average time series in each of the 400 cortical regions. Functional connectivity matrices are then obtained via computation of the correlation coefficient of time series for each pair of regions, followed by a Fisher r to z transformation. The result of this process is a 400 × 400 RSFC matrix for each individual.

### 2.4 Computing microstructure-function coupling coefficients

Microstructure-function coupling coefficients are computed on a per-subject, per-region basis via comparison of a given region’s MPC and RSFC features. Specifically, we compute the Pearson correlation between each row of a subject’s MPC matrix with the corresponding row of the same subject’s RSFC matrix. This results in a 400 × 1 vector of coupling coefficients for each subject. A high coupling coefficient indicates that a region is functionally connected to regions with similar microstructure and vice versa.

### 2.5 Identifying coupling components with NMF

To identify regional patterns of structure-function coupling and associated inter-individual variability, we employ non-negative matrix factorization (NMF). NMF is a matrix decomposition technique which seeks to decompose an input matrix into a set of additive components (D. D. Lee & Seung, 1999; D. Lee & Seung, 2001) (see Supplementary Materials, Section 1.3 for details). In our implementation, the input data matrix is a *m x s* matrix, with each m row being a brain region, and each s column representing a given subjects coupling regional coupling coefficients (*m* = 400, *s* = 384). To construct the input matrix, each subject’s 400 × 1 array of coupling coefficients (**Section 2.5**) is stacked side by side to form a 400 × 384 matrix. This matrix is then shifted by its minimum value to satisfy the non-negativity requirements of NMF while maintaining its distribution. The resulting shifted matrix is input to NMF. We implemented NMF using the sklearn package (version 0.20.3) and used a non-negative double singular value decomposition initiation procedure, which promotes sparsity in the NMF outputs (Boutsidis & Gallopoulos, 2008) to aid in the final interpretation of the resulting neuroanatomical patterns (Patel, Steele, Chen, Patel, Devenyi, Germann, Tardif, & Chakravarty, 2020; Sotiras et al., 2015). While we do promote sparsity via this initialization procedure, notably we employ NMF without any orthogonality constraint unlike some recent applications (Nassar et al., 2018; Patel, Steele, Chen, Patel, Devenyi, Germann, Tardif, & Chakravarty, 2020; Sotiras et al., 2015, 2017). This is done in order to allow overlapping patterns of coupling variability as opposed to enforcing each brain region to occupy only one of the output components/patterns, as we have done in other recent work (Patel et al., 2022). We select the optimal number of components to analyze, *k*, based on a previously developed stability analysis (Patel, Steele, Chen, Patel, Devenyi, Germann, Tardif, & Chakravarty, 2020) which balances the accuracy of the decomposition, assessed via gradient of the reconstruction error, with the spatial consistency of the identified components, assessed via a stability coefficient, across varying subsets of participants.

#### 2.5.1 Interpreting NMF output

An NMF decomposition contains two outputs - the *W* component matrix and *H* weight matrix. *W* contains the component scores of each brain region, which can be mapped back to the cortical surface in order to visualize the spatial location of a given component. To aid in the interpretation of the output patterns in terms of cytoarchitectonic types, we apply a cortical type atlas (Mesulam, 2000; Paquola et al., 2019; Valk et al., 2022) in order to identify the cytoarchitectonic distribution of component scores. If a subject has a high weighting (*H* value) for a given component, this is indicative of a high coupling coefficient in the brain regions occupied by said component. In this way, the combination of *W* and *H* describes the additive parts of the original coupling data, and can be used to identify spatial and inter-individual patterns of coupling covariation.

### 2.6 Identifying the cognitive determinants of Structure-Function Coupling

Using the NMF derived inter-individual coupling variability, we next sought to assess how variability in structure-function maps onto cognition. To do so we capitalized on the in depth cognitive phenotyping conducted by the HCP and included a number of tests across a range of cognitive functions including: episodic memory, cognitive flexibility, inhibition, fluid intelligence, language decoding, vocabulary comprehension, processing speed, impulsivity, spatial orientation, sustained attention, verbal episodic memory, and working memory. Specific instruments used to assess these tasks can be found in **Table S1** as well as in (Barch et al., 2013). Note that, where possible, we used age adjusted measurements for each of the tests mentioned above.

#### 2.6.1 Deriving latent variables related to microstructure-function coupling and cognition: Partial Least Squares Analysis

To investigate the relationship between NMF derived variability in structure-function coupling and behaviour, we employ PLS correlation analysis (Krishnan et al., 2011; McIntosh & Lobaugh, 2004; McIntosh & Mišić, 2013). PLS is a multivariate technique which seeks to optimize the covariance between two datasets. In doing so, PLS computes latent variables (LV), each of which contain a set of weighting variables that describe covarying relationships between the two input datasets. Here, the two input datasets are NMF subject weights from the *H* weight matrix (brain data), and performance across the cognitive tests described above (behaviour data).

Thus, we use PLS to identify LVs which describe linear combinations of the NMF weights and cognitive performance that maximally covary, describing multivariate mappings between inter-individual variability in structure-function coupling and cognitive performance. Permutation testing was used to assess the significance of each latent variable and bootstrapping was used to assess the reliability of the contributions of each of the variables (see Supplementary Materials, Section 1.4 for further details).

### 2.7 Transcriptomic analyses

To further characterize our NMF-derived components, we performed a cell-type enrichment analysis to identify unique and overlapping cell-type expression patterns of the output components (Seidlitz et al., 2020). Data from six post-mortem adult brains (Allen Human Brain Atlas, AHBA) were preprocessed using the abagen toolbox (Markello et al., 2021) to obtain a 6932 (samples) x 15633 (genes) expression matrix describing gene expression patterns at different locations. We then applied spatial component maps to the expression data in order to obtain a samples x genes matrix describing gene expression values of samples located within a given component.

To identify gene expression patterns unique to NMF components, we used mean centered PLS (mcPLS). mcPLS searches for orthogonal latent variables which express covarying patterns of two input data matrices X and Y (see Supplementary Material, Section 1.5). Statistical significance of LVs and stability of gene weights are obtained via permutation testing and bootstrap resampling, respectively, similar to behavioural PLS. For each significant LV, we obtained a positive (PLS+) and negative (PLS-) gene set, each containing those genes with a BSR higher/lower than 2.58/-2.58. These gene sets are determined to describe expression patterns separating components and are thus further analyzed.

#### 2.7.1 Cell-type enrichment

We performed a cell-type enrichment analysis to identify cell classes preferentially represented by the genes in the PLS+ and PLS- sets computed by mcPLS (Hansen et al., 2021; Seidlitz et al., 2020). Clustering of the spatial patterns of 58 cell types in the Allen Human Brain Atlas has identified seven cell classes: astrocytes, endothelial cells, microglia, excitatory neurons, inhibitory neurons, oligodendrocytes and oligodendrocyte precursor cells (Seidlitz et al., 2020). We computed the ratio of genes in each of the PLS+ and PLS- gene sets that are preferentially expressed across each cell type, as in previous work (Hansen et al., 2021; Seidlitz et al., 2020). Significance was assessed via permutation testing, whereby enrichment ratio distributions were computed via 10,000 repetitions each drawn using a random set of genes.

## 3.0 Results

### 3.1 Data

384 unrelated participants from the HCP Young Adult Dataset were selected (mean age = 28.6 +- 3.77, F/M = 202/182, Handedness = 67.2 +- 40.58).

### 3.2 Structure-Function coupling follows cytoarchitectonic type patterns

At the group level, we found that the link between microstructural similarity, assessed via MPC, and connectivity, assessed via RSFC, displayed a regionally heterogeneous pattern with strong coupling in primary cortices and weaker coupling in transmodal cortices, in line with previous findings (Huntenburg et al., 2017, 2018; Paquola et al., 2019; Valk et al., 2022). For each individual, we performed a row-wise correlation of their MPC and RSFC matrices to obtain a coupling coefficient for each of 400 brain regions. These values are then averaged across subjects to obtain the mean regional coupling coefficients displayed in **Figure 2A**. While visually apparent that mean coupling is highest in primary motor and visual cortices and lowest in temporal, frontal and parietal cortices, we apply a cytoarchitectonic classification atlas (Mesulam, 2000) to this mapping to quantify the differences in mean coupling across cortical types (**Figure 2A**). This confirms that structure-function coupling is strongest in idiotypic cortices including primary motor and visual areas (mean coupling coefficient = 0.32 +/-0.11). Unimodal (mean = 0.11 +/- 0.12) and heteromodal (mean = 0.05 +/- 0.08) cortices display comparable coupling but markedly lower than idiotypic regions. Limbic cortices (mean = 0.02 +/- 0.07) show the lowest mean coupling, though the decrease in coupling from heteromodal to limbic is much less than from idiotypic to heteromodal/unimodal. We also plotted the node wise variability in coupling measurements in **Figure 2B** by, for each region, computing the standard deviation of coupling coefficients across all subjects. Similar to mean coupling, we display a grouping of coupling variability by cytoarchitectonic class. Here, cortical types are much less differentiated than the mean coupling values, suggesting that individual variability in regional coupling is not specific to cortical type.

**Figure 2.**
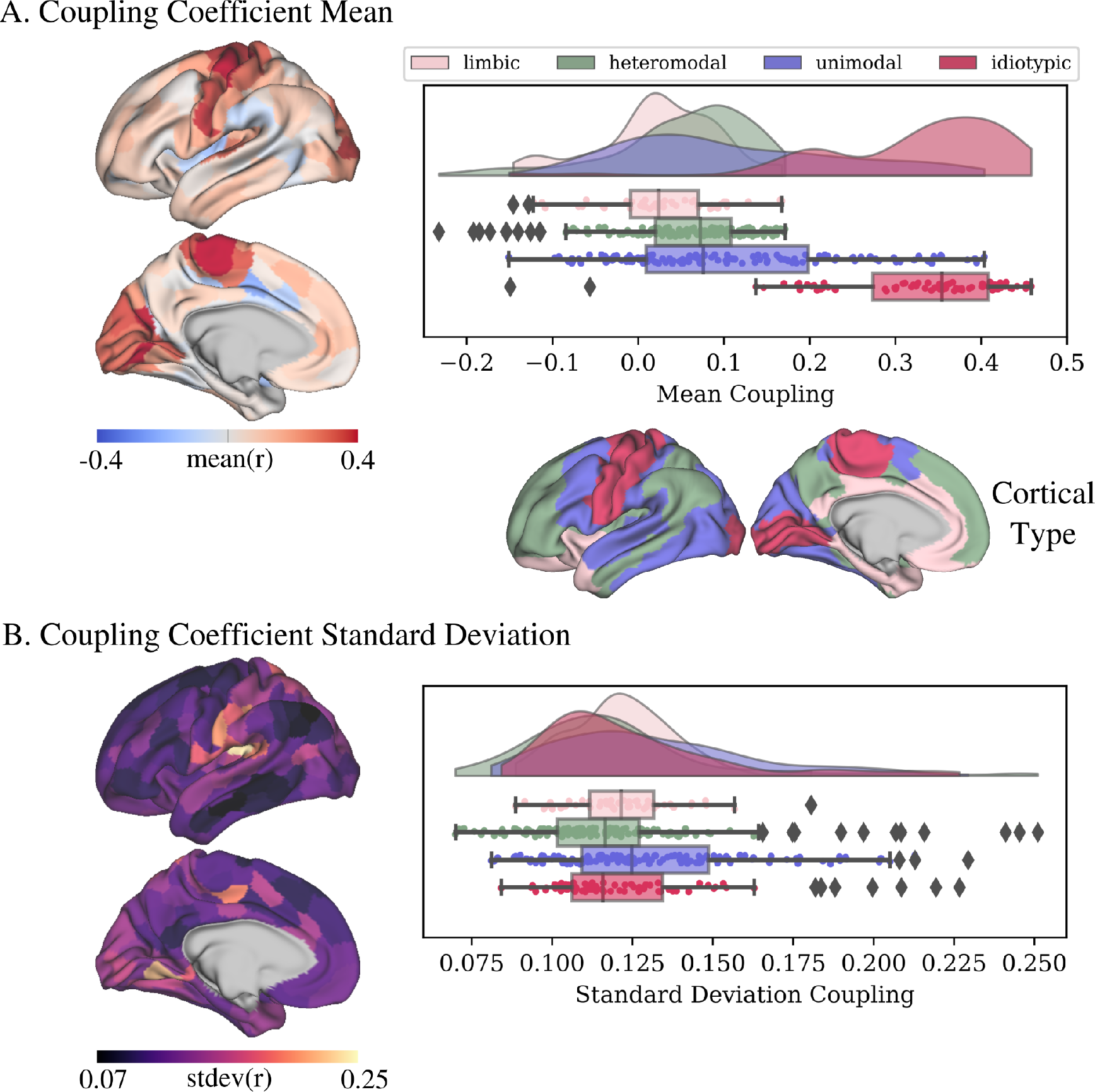
**Microstructure-function coupling varies across the cortex**. The relationship between microstructural similarity and resting state functional connectivity varies markedly across the cortex. Panel A plots the mean coupling coefficient, panel B the standard deviation of each region across subjects. The cortical type atlas is colour coded according to cytoarchitectonic type. (A) In idiotypic cortices including primary motor and visual regions, coupling is strongest. Conversely, the link between structure in function drops off notably in heteromodal, unimodal, and limbic regions. (B) No specific cortical type shows markedly different variability compared to others, when looking at the regional standard deviation of coupling across all subjects.

### 3.3 NMF Identifies Four Coupling Components

Using a previously established stability analysis (Patel, Steele, Chen, Patel, Devenyi, Germann, Tardif, & Chakravarty, 2020), we identified four components as a suitable balance between accuracy and consistency across subjects. Briefly, subjects are split into two groups and NMF is run separately for each group. The spatial similarity of the output components is measured between the two groups (stability coefficient) and the accuracy of the decomposition (reconstruction error) is also assessed. This process is repeated for 10 splits of subjects at each granularity, *k*, from 2 to 40. **Figure S1** plots the stability coefficient and change in reconstruction error across this range. Stability is highest at k=2, decreases until k=6 and then levels off. Meanwhile, the large changes in reconstruction error at *k*=3 and *k*=4 convey that at *k*<=4, major patterns of the input data are yet to be captured. Based on this, we select *k*=4 for further analysis.

### 3.4 Coupling Components Match Cortical Type Transitions

The 4 component solution is displayed in **Figure 3**. Each column of the *W* matrix is mapped back to the cortical surface to display the spatial location of each component (**Figure 3A**). Each component captures a pattern of coupling covariation. Here we discuss the anatomical patterns identified by each component. In order to better describe, and compare the anatomical features of each component, we apply a cytoarchitectonic atlas (Mesulam, 2000) to each component score map to derive raincloud plots which show the distribution of component scores within each cortical type (**Figure 3A**). To aid in the comparison of component spatial patterns, we rank regions based on their component scores and subtract the resulting rankings for each pair of components (**Figure 3B-G**). Specifically, for each component, we rank regions from 0 (lowest component score) to 400 (highest component score), then subtract these rankings from each other. A value of 400 indicates that a given region ranked the highest in one component and the lowest in another, and thus one component is much more prominent in said region than the other. Conversely, a value of 0 indicates the region is of equal importance to each component.

**Figure 3:**
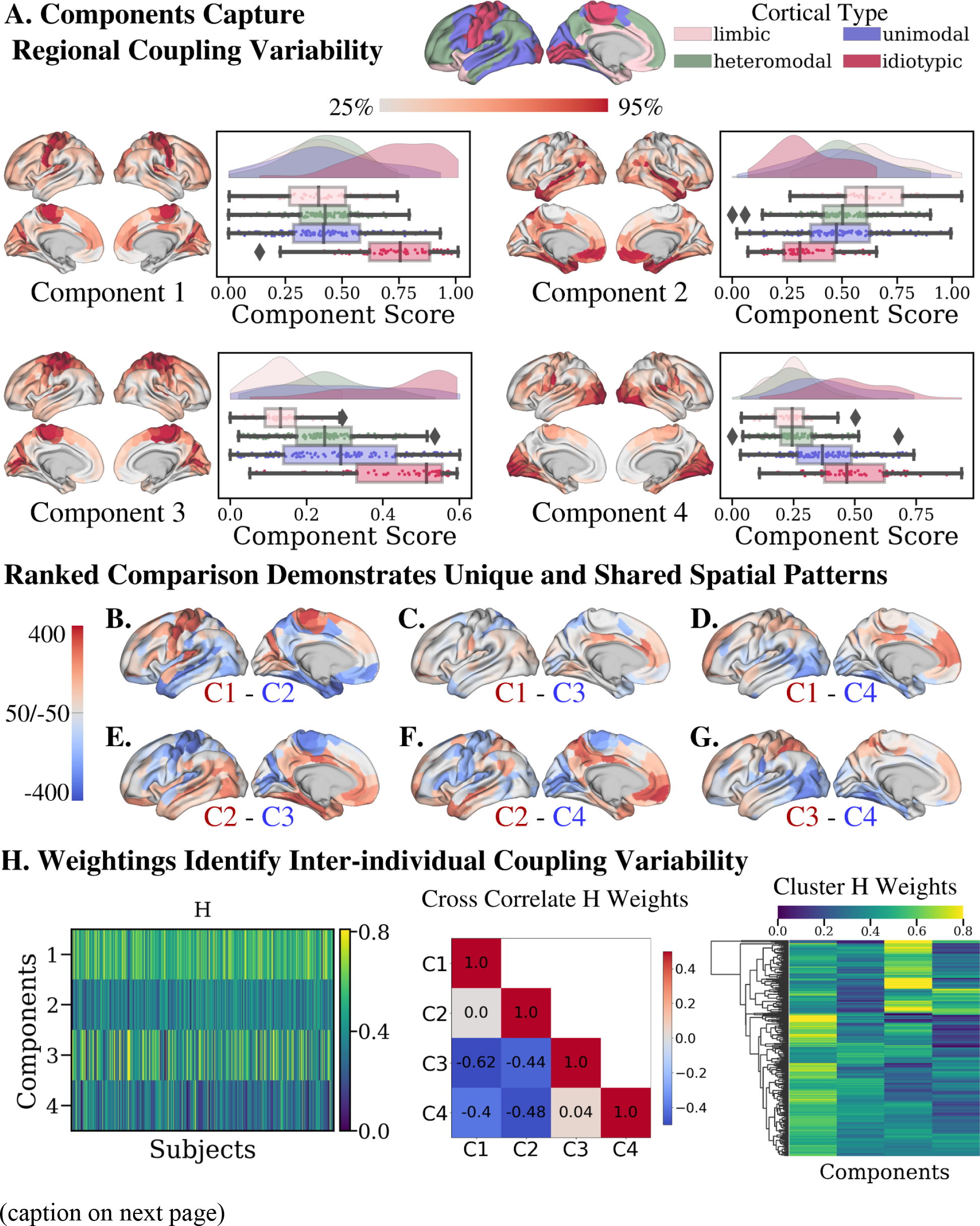
**Coupling components capture both regional and inter-individual variability**. (A) Mappings of each of the 4 components. Left and right hemispheres are shown, colour coded by their component score. The range of each component varies, so each is colour coded according to their own range of 25% to 95%, such that dark red represents the 9th percentile of each component. Raincloud plots for each component show the distribution of component scores within each of 4 cytoarchitectonic classes. The Cortical Type atlas at the top of the figure displays these classes for reference. (B-G) Ranked comparisons of spatial components. For each component, we rank each of 400 cortical regions based on component score (0=lowest, 400=highest). Then, for each pair of components we subtract rankings, and map results back to the cortical surface. For each map, the order of subtraction is written below. For example, for the C1-C2 map, a value of 400 (red) indicates a region that has the highest C1 score but the lowest C2 score. (H) Weight matrix of the 4 component decomposition is displayed on the left, where each row is a component and each column a subject. A cross correlation matrix displays the relationship between component weights, allowing inference of how the components separate individuals. On the right, a hierarchical clustering is presented for visualization purposes

Component 1 occupies idiotypic cortices. Specifically, the highest scores (dark red) can be found throughout the motor cortices, with high values visible in visual areas as well. However, component 1 is not solely confined to these regions, as moderate values (light red) can be found throughout lateral and medial frontal regions, as well as some portions of the parietal cortex. This is reflected in the cytoarchitectonic distribution plots which shows prominently higher scores in idiotypic cortex (red), but moderate and comparable values in unimodal (blue), heteromodal (green) and limbic (pink) cortices. Interestingly, we find that the spatial pattern of component 1 resembles the group average coupling coefficient map in **Figure 2**, with high values in primary motor and visual regions, low to moderate coupling in widespread frontal, temporal, and parietal areas, and low/decoupling in regions such as the insula, tempo-parietal junction, and select medial frontal regions.

Component 2 displays a markedly different pattern compared to component 1 (**Figure 3B**). This component is most prominent in limbic cortices, with high component scores (dark red) occupying the bilateral medial temporal lobe, temporal pole, and medial orbitofrontal cortex. This component also occupies heteromodal regions strongly, with high scores in the middle temporal gyrus, and moderate (light red) scores found in many other regions, including the medial parietal and frontal regions, and select regions of the inferior parietal lobe. Lower values are then found in the unimodal cortices, with the lowest in the idiotypic cortex. Unlike component 1, more separation can be seen between limbic, unimodal, and heteromodal regions such that the cytoarchitectonic distribution plots show a gradual transition in component 2 scores from limbic (highest) to heteromodal, unimodal, then idiotypic (lowest).

At first glance, component 3 looks highly similar to component 1 in terms of its spatial pattern. Indeed, component 3 is clearly prominent (dark red) in idiotypic cortex such as primary motor and visual cortex (**Figure 3A)**. Unlike component 1 however, limbic cortices display very low component scores, much lower than that of heteromodal and unimodal cortices. Another unique feature of this component is the distribution of component scores in unimodal regions. In the cytoarchitectonic distribution plot, unimodal cortices (blue) display a near uniform distribution across the entire range, in contrast to the more normally shaped distributions shown by unimodal cortex in other components. Thus, component 3 is marked by strong separation of idiotypic and limbic cortices, with other regions displaying a much more variable pattern.

Component 4 is most prominent in bilateral visual cortex, and also displays moderate scores in motor regions. Indeed, this is evident in the higher values in idiotypic regions displayed in the cytoarchitectonic distribution plot. Component 4 also includes moderate scores in regions classified as unimodal cortex such as the inferior temporal lobe. Heteromodal and limbic cortices display similarly low scores in component 4. While similar to components 1 and 3 in terms of the idiotypic preference, component 4 displays a stronger localization to visual cortices as opposed to motor, or motor and visual regions (**Figure 3B**). Indeed, motor cortex in component 4 is spanned by moderate light red values as opposed to the dark red evident in the visual cortex.

### 3.5 NMF Weightings Demonstrate Inter-individual Coupling Variability

In **Figure 3H** we plot the *H* weighting matrix from the 4 component solution. This matrix has 1 row for each component, and one column for each subject. Thus, *H(:,j)* is a vector describing how subject *j* loads onto each component. Recalling that NMF computes *W* and *H* such that *X* ≈ *WH*, a high weighting for component 1 suggests that, through additive reconstruction of the input data, a given subject has high structure-function coupling in the regions occupied by component 1. Thus, we use *H* to assess the structure-function coupling of each individual within each of the spatial patterns described in **Section 3.4**.

To further probe patterns of inter-individual variability we create a cross correlation matrix of *H*, that is we correlate the weightings in each pair of components (**Figure 3H**). Here we see that there are some relations between component weightings. Specifically, component 1 is inversely correlated with components 3 and 4. Thus, an individual with high weighting in component 1 will, through additive reconstruction, express coupling in component 1 regions but less so in components 3 and 4. We find this inverse relationship interesting in light of the previously discussed spatial patterns. As indicated by the grey colouring in **Figure 3C**, both components 1 and 3 are prominent in the idiotypic cortices, thus regardless of an individual’s weighting pattern the expected strong structure-function coupling in idiotypic cortex is realized. Where they differ is in the separation of idiotypic and limbic regions. As indicated by the light red regions in limbic cortices in **Figure 3C**, individuals who obtain their idiotypic coupling via high component 1 weight will have increased coupling in limbic regions compared to those who obtain it via a high component 3 weighting. Similarly, component 1 weightings are inversely correlated with component 4 weightings and component 4 displays a similar, though not as marked, separation of limbic and idiotypic regions in its cytoarchitectonic distribution plot (**Figure 3D**). These comparisons suggest that while structure-function coupling in idiotypic regions is consistently high, there exists variability in limbic cortices such that a subset of the individuals at study display increased coupling. Conversely, other individuals show decreased coupling in limbic cortices and are characterized by a much stronger separation in degree of coupling across cortical types.

This is furthered via consideration of component 2, which is most prominent in limbic regions, and displays a gradual decrease to heteromodal, unimodal, and idiotypic cortices. The cross correlation matrix tells us that, like component 1, individuals who load high onto limbic heavy component 2 are expected to show inversely low weightings onto component 3 and 4. This furthers the idea of a subset of individuals who display increased coupling in limbic regions. To supplement these results, we present a hierarchical clustering of the *H* matrix in **Figure 3H** as well. While not presented as a formal analysis, grouping subjects according to their weighting patterns allows a clear visualization of certain groups of individuals showing inverse and defined weighting patterns (**Figure 3H**). When considering our results in the context of previously established group level structure-function coupling patterns (Huntenburg et al., 2017), these results suggest that non-idiotypic cortices which, on average, display low coupling do indeed exhibit some degree of inter-individual variability.

### 3.6 Behavioural relevance of coupling variability

To assess the behavioural relevance of the identified structure-function coupling variability, we next ran a PLS brain-behaviour analysis with the brain data being a subjects x components matrix, where each column denotes an individual’s *H* weight, and the behaviour data being a subjects x behaviours matrix where each column denotes performance on one of 19 cognitive tests assessed by the HCP. PLS analysis output one significant LV (p=0.008) which explained 72.7% of covariance between brain and behaviour. **Figure 4A** shows a scree plot displaying the pvalues and covariance explained for each of the identified LVs. As described in **Section 2.6.1**, to ascertain the variables involved in an LV we apply a bootstrap ratio (BSR) threshold of 2.58 for brain variables, and discuss only behaviour variables for which a 95% confidence interval (CI) does not overlap 0. Using these criteria, the brain variables involved in LV1 are increased weights in components 1 and 2, and decreased weightings in component 3 (**Figure 4B**). LV1 associates this pattern with a behavioural pattern of increased performance on tasks of episodic memory (r = 0.11, CI = [0.05,0.11]), cognitive flexibility (r = 0.07, CI = [0.017,0.073]), inhibition (r = 0.14, CI = [0.019,0.18]), vocabulary comprehension (r = 0.12, CI = [0.02,0.14]), processing speed (r = 0.14, CI = [0.02,0.19]), delay discounting (impulsivity) task via both the 200 (r = 0.11, CI = [0.08,0.15]) and 40K (r = 0.13, CI = [0.03,0.2]) AUC measures, increased correct trials (r = 0.07, CI = [0.04,0.17]) but longer reaction times (r = 0.16, CI = [0.13,0.22]) in the spatial orientation task, and increased performance in the sustained attention task specificity (r = 0.07, CI = [0.03,0.13]) (**Figure 4C**). Thus, this LV describes an overall pattern in which individuals weighing heavily onto components 1 and 2, but less so on component 3, display a general pattern of increased cognitive performance across a range of cognitive tasks and domains.

**Figure 4:**
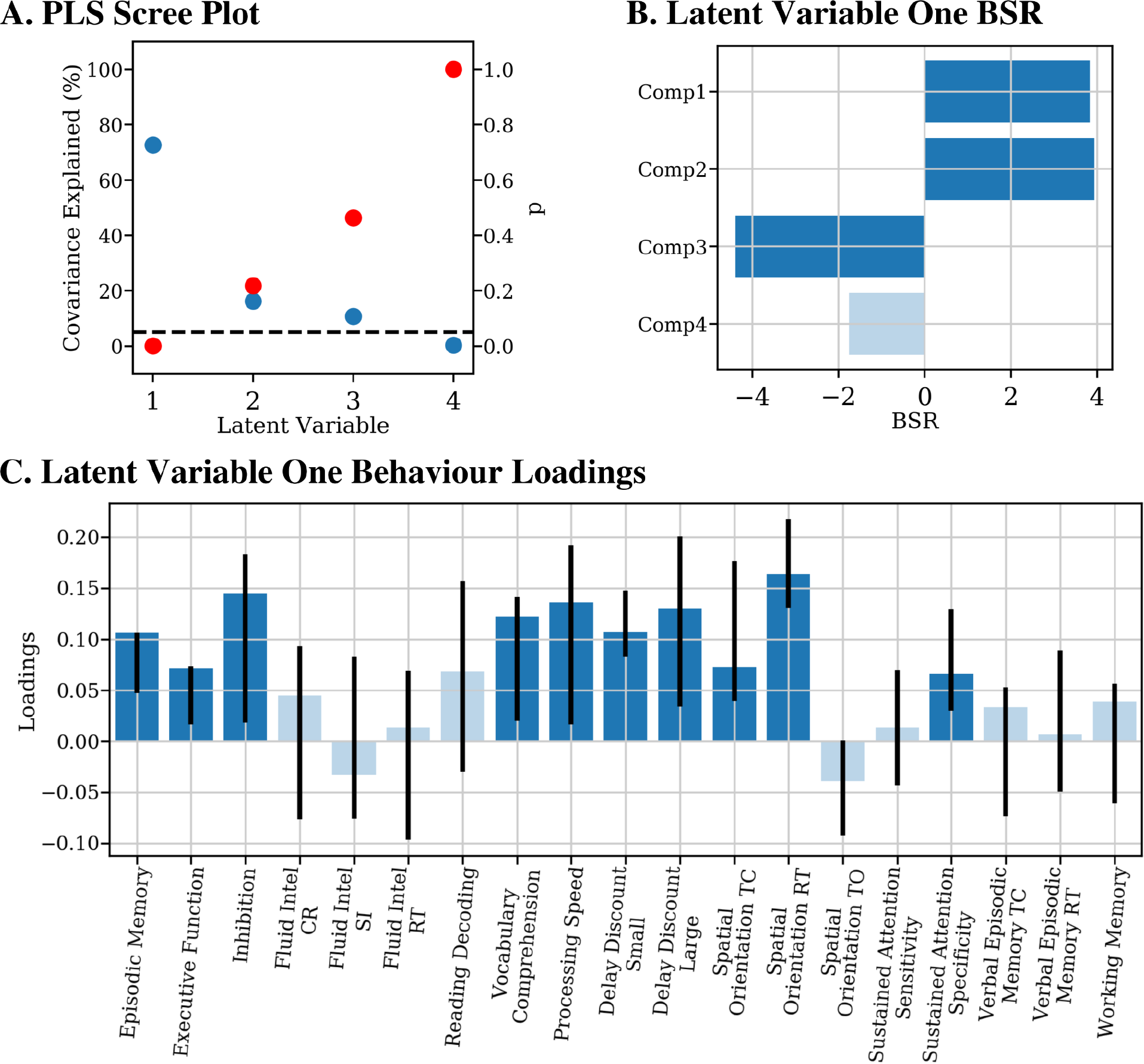
PLS analysis identifies one coupling-behaviour latent variable. (A) Scree plot shows, for each LV, the covariance explained (blue) and non parametric p value (red) obtained via permutation testing. A dotted line is drawn at p=0.05. Only LV1 is significant at this level. (B) Bar plot showing the BSR of each component. Solid blue indicates components for which their BSR was above a threshold of 2.58. (C) Bar plot showing the loading of each behaviour variable within LV1. The bar is drawn to the height of a given variable’s loading, while the error bars indicate the range of the 95% confidence interval computed via bootstrapping. Variables for which the 95% CI crosses zero appear transparent, compared to the solid blue bars indicating a 95% CI that does not cross zero.

In **Section 3.4** we presented NMF results suggesting that, while all individuals display high coupling in idiotypic regions, the way in which our coupling input data is reconstructed via the 4 components varies. Specifically, a subset of individuals were found to show a degree of increased structure-function coupling in limbic cortices, in addition to the expected coupling of idiotypic regions. In this context, the covariation of increased component 1 and 2 weightings, and decreased component 3 weightings with a general pattern of increased cognitive performance provides a potential cognitive determinant of this NMF derived coupling variability.

### 3.7 Cell-type enrichment of components

To further characterize the obtained components, we used mcPLS to identify gene sets (PLS+ and PLS-) describing component-specific expression patterns, and then identified canonical cell classes represented by the PLS+ and PLS- gene sets. Our mcPLS analysis identified three significant latent variables, with discussion here focussed on the first LV (variance explained = 70%, p<0.01). LV1 shows that component 1 and 4 had common gene expression patterns, a pattern which was in contrast to the expression pattern associated with component 2 (Figure 6A). Thus, similar to the cortical type analysis described in Section 3.4, LV1 contrasts idiotypic (components 1 and 4) and limbic (component 2) cortices, here in terms of gene expression. Cell-type analysis showed that PLS+ genes (with increased expression in idiotypic components 1 and 4) are enriched for endothelial cells, as well as excitatory and inhibitory neurons. Meanwhile, PLS- genes (with increased expression in limbic component 2) were enriched for astrocytes, microglia and oligodendrocyte precursor cells.

**Figure 6:**
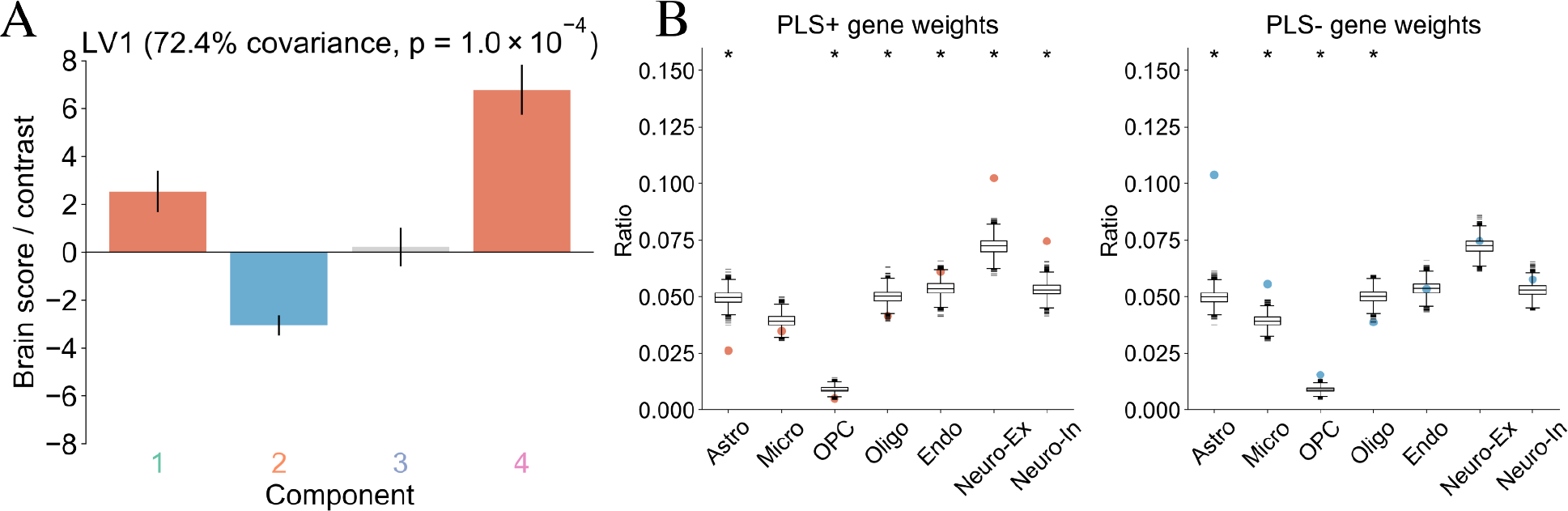
**Cell type enrichment analysis further separates idiotypic and limbic cortices**. (A) Contrast plot showing the extent to which each NMF component is associated with either the positive (PLS+) or negative (PLS-) weighted genes of LV1. Error bars denote the 95% confidence interval in LV1 scores for each component. LV1 captures differences between components 1 and 4 vs. component 2. (B) Ratios of genes in PLS+/PLS- gene sets preferentially expressed in 7 cell types. Coloured dots (red, blue) represent the enrichment ratio, while box plots show the null distribution of enrichment ratios derived from a sample of 10,000 random gene lists.

## 4.0 Discussion

In this work we study inter-individual variability of microstructural-functional coupling in the human cortex. We use NMF to identify four coupling components which decompose group-level data into regional patterns of variability, and quantify individual differences in microstural-functional coupling. Using a multivariate approach, we then show that specific patterns of coupling map onto a latent factor of increased performance across a range of cognitive domains. We also show that the identified coupling components map to variations in cell type.

### 4.1 Structure-Function coupling maps to cortical and cell type transitions

We begin our analysis by replicating and extending previous group level findings on the spatial heterogeneity of structure-function coupling patterns. We show that microstructural similarity, assessed via MPC, and connectivity, assessed via RSFC, displayed a regionally heterogeneous pattern with strong coupling in primary cortices and weaker coupling in transmodal cortices. Specifically, we used NMF to identify four coupling components and applied a cytoarchitectonic atlas (Mesulam, 2000) to confirm that structure-function coupling is strongest in idiotypic cortices, followed by unimodal, heteromodal, and finally limbic regions. This is in line with previous works that have identified a similar cortical heterogeneity using a variety of data types (Huntenburg et al., 2017, 2018; Paquola et al., 2019; Suárez et al., 2020; Vázquez-Rodríguez et al., 2019). We further the characterization of these cortical differences by performing a cell-type analysis. Here, we show that sets of genes preferentially expressed within the idiotypic cortices are more enriched for neurons (inhibitory neurons, excitatory neurons, endothelial cells), while gene sets preferentially expressed in limbic cortices are instead more enriched for neuron support (astrocytes and microglia). Thus, we extend previous findings to show that at the group level, heterogeneity in structure-function maps not only to differences in cortical types but also cell type.

### 4.2 Coupling Components show Variability in Transmodal Regions

As the majority of previous studies have focussed on structure-function coupling at the group level (Suárez et al., 2020; Sydnor et al., 2021), a key goal of the present work is to study the existence of inter-individual variability in microstructural-functional coupling in transmodal cortices. An unexplored question in this context, and in the broader field of structure-function analysis, is the influence of individual variability on group level findings of decreased coupling in limbic regions (Suárez et al., 2020).

Three of the four NMF components identified regional coupling patterns in line with established work. Components one, three, and four show a high degree of spatial overlap (**Figure 3C, 4D, 4G**), and are all most prominent in idiotypic cortex such as primary motor and visual areas. The cross-correlation of component weightings demonstrate that individuals loading heavily onto component one show an inverse pattern for components three and four, which has two significant implications. First, this confirms that the majority of individuals do show strong coupling in these regions, as those who do not load heavily onto component one will instead load onto components three and four. Via additive reconstruction of component one, or components three and four, NMF decomposition shows most individuals with high coupling values in these regions. This is in line with a number of relevant previous studies, including those studying microstructure-function relationships (Huntenburg et al., 2017; Paquola et al., 2019; Valk et al., 2022), but also those studying structure-function relationships via diffusion MRI estimates of structural connectivity (Gu et al., 2021; Mišić et al., 2016; Preti & Van De Ville, 2019; Suárez et al., 2020; Vázquez-Rodríguez et al., 2019), those using partial correlation based estimates of resting state connectivity (Gu et al., 2021; Liégeois et al., 2020), and even those employing task-based fMRI estimates of functional connectivity (Wu et al., 2020). Thus, our findings support a growing body of work, across a range of methods and metrics, that support delineations between primary and transmodal cortical regions based on the underlying cytoarchitectonic, myeloarchitectonic, and functional properties.

Second, the inverse correlation of these seemingly overlapping components begs the question of why an individual may weigh onto one but not the other component, and what their differences are. Indeed, while spatial patterns are similar, some important differences are noted. To aid in this interpretation, we applied a cytoarchitectonic atlas (Mesulam, 2000; Paquola et al., 2019; Valk et al., 2022) onto each spatial component to identify the distribution of component scores, and thus prominence of each component, within each of four cortical types. These cytoarchitectonic distribution plots show that, while each component is prominent in idiotypic cortex, components three and four separate idiotypic and limbic cortices much more strongly than component one, which instead is more evenly distributed across limbic, heteromodal, and unimodal cortex (**Figure 3A, 4C, 4D**). Thus, while all individuals demonstrate strong coupling in idiotypic cortex, the accompanying coupling patterns in transmodal regions vary depending on an individual’s expression of the identified components. This differential loading is indicative of coupling variability in limbic, unimodal, and heteromodal regions, but not in idiotypic cortex where coupling is consistently strong.

Visual inspection of component 2 makes clear that the spatial pattern is unique relative to other components. Indeed, the cytoarchitectonic distribution plot of component 2 shows that it is, in sharp contrast to components 1, 3, and 4, most prominent in limbic cortices, with transitional decreases to heteromodal, unimodal, and finally idiotypic cortex respectively. Therefore, when comparing individuals with low vs. high weighting, component two does not capture differences in idiotypic coupling, but instead separates individuals with variable microstructural-functional coupling in limbic and heteromodal regions. This finding suggests that low group-level coupling measures in non-idiotypic cortex are not solely driven by consistently low coupling measurements but instead a degree of variable microstructure-function relationships across individuals. The unique spatial pattern of component 2, in comparison to other components and group average coupling, is also of interest in the context of previous studies of cortical microstructure and functional variation. Cortical gradients of microstructure have been identified showing transitions from primary cortices to limbic regions (Huntenburg et al., 2017, 2018; Paquola et al., 2019). Meanwhile, a number of studies have investigated variations in resting state based fMRI patterns to identify functional hierarchies with primary regions at one end, and heteromodal regions including the default mode network at another (Margulies et al., 2016; Paquola et al., 2019). While components 1, 3 and 4 show overlap with expected coupling patterns, component two is instead most prominent in these areas of divergence including limbic and heteromodal regions. Thus, while previous studies establish that microstructural and functional variation show divergence in transmodal regions, our findings suggest that the degree of divergence varies at the individual level. Interestingly, cross correlation of component weightings shows that, like component one, individuals with high component two weights have lower weights for each of components three and four. This furthers the idea that the identified components and weightings can be used to describe coupling variability in transmodal regions. Interestingly, studies investigating structure-function using diffusion weighted imaging derived measures of structural connectivity display a similar finding (Mišić et al., 2016; Preti & Van De Ville, 2019; Suárez et al., 2020; Vázquez-Rodríguez et al., 2019). Thus, this regional variation in structure-function coupling which separates primary regions from transmodal areas is a consistent finding across methodologies and metrics. Apart from the definition of primary and transmodal areas via microstructure-function coupling, there is further correspondence with gene expression profiles and functional processing hierarchies (Huntenburg et al., 2018). The basis of decoupling in transmodal cortices remains unclear. These regions have expanded evolutionarily (Hill et al., 2010; Reardon et al., 2018; Smaers et al., 2017; Toro et al., 2008), in comparison to non human primates and other species which formed the basis of the theory of microstructural similarity as a connectivity principle. Thus, it has been suggested that these evolutionary changes are accompanied by fewer constraints of structure on function which enable a more variable connectivity pattern (Buckner & Krienen, 2013).

### 4.2 Cognitive associations of coupling variability

Using our PLS analysis, the NMF components, and a broad range of cognitive domains we intentionally used a hypothesis free approach to understand this relationship The LV identified related a cognitive mapping of the previously discussed separation of regional coupling patterns, such that individuals with increased microstructural-functional coupling in idiotypic but also limbic and, to a lesser degree, heteromodal and unimodal cortices perform better on nearly all cognitive domains assessed. Strong coupling in primary motor and visual regions is consistent across individuals, but also across species (Valk et al., 2022), suggesting a behavioural and/or cognitive benefit to this conserved feature. Conversely, decoupling in transmodal regions has been suggested as a possible evolutionary development whereby relaxed constraints enable increased plasticity (Buckner & Krienen, 2013; Valk et al., 2022). Our PLS results however show that component 2 weightings, and therefore coupling in limbic and heteromodal regions, is associated with increased cognitive performance. Thus, these results suggest that some degree of increased coupling may be beneficial in the context of cognition.

Comparison to previous findings is challenging as, to our knowledge, this is the first work specifically testing microstructural-functional variability in relation to cognitive performance. This has been tested previously in studies which employed diffusion MRI to measure structural connectivity as opposed to microstructural similarity. For example, Medaglia and colleagues found that the degree of alignment between functional and anatomical networks supported variation in cognitive flexibility (Medaglia et al., 2018). Structure-function relationships have also been shown to be heavily influenced by development, such that developmental changes in structure-function impact executive function abilities (Baum et al., 2020). These findings suggest that individual level variations in structure-function, assessed via diffusion MRI, may impact cognition. Direct relevance to our study, which employs different measurements, is cautioned but may suggest that a cognitive determinant of microstructural-function coupling variability is also plausible.

### 4.3 Studying microstructure-function relationships in vivo

The study of cortical microstructure has classically been conducted via post-mortem analysis of cyto- and myelo-architectonics, identifying spatial variations in the laminar structure of the cortical sheet (Brodmann, 1909; Palomero-Gallagher & Zilles, 2019; Vogt & Vogt, 1919). In this study, we use the ratio of T1w/T2w as a microstructural marker. This marker was originally proposed as an index of cortical myelin, due to correspondence between cortical T1w/T2w patterns and known variations in cortical myelin (Glasser et al., 2014, 2016; Glasser & Van Essen, 2011). However, based on the non quantitative nature of both T1w and T2w (Tardif et al., 2016; Zatorre et al., 2012), specificity of T1w/T2w to myelin has been questioned. Indeed, the degree of correlation between T1w/T2w and quantitative markers of myelin has been shown to vary (Arshad et al., 2017; Prasloski et al., 2012; Righart et al., 2017; Uddin et al., 2018, 2019). Thus, if specificity to cortical myelin was desired, quantitative metrics such as myelin water fraction (Prasloski et al., 2012) or quantitative T1 (Huntenburg et al., 2017; Waehnert et al., 2016) may be more desirable. However, that T1w/T2w captures unique components of microstructure has also been demonstrated (Arshad et al., 2017; Prasloski et al., 2012; Righart et al., 2017) which supports the use of T1w/T2w as a microstructural index. Importantly, a recent study showed that cortical T1w/T2w measurements closely matched systemic variations in cortical lamination patterns (John et al., 2021), which has direct relevance to its use in our study. Here, we measure microstructural similarity via the recently developed MPC (Paquola et al., 2019). In this approach the T1w/T2w signal is sampled along multiple cortical depths to create a microstructural profile at each point along the cortical surface, and pairwise comparisons of microstructural profiles indicate how similar two cortical points are according to their depth dependent measurement of T1w/T2w signal (Paquola et al., 2019). Crucially, this approach follows the spirit of classical post-mortem analysis by querying depth-varying information and has indeed been shown to recapitulate known variations in the laminar structure of the cortex (Paquola et al., 2019).

According to the structural model (Barbas, 2015; Barbas & Rempel-Clower, 1997; Beul et al., 2017; García-Cabezas et al., 2019) microstructural similarity is indeed indicative of the connectivity of two regions, a finding generated based largely on animal models (Pandya et al., 2015), but also supported by findings in humans (van den Heuvel et al., 2016). This, as well as the overlap in regional coupling of microstructure and structural connectivity studies, suggests that both MRI derived approaches are querying the same phenomenon of brain organization albeit in slightly different ways. However, they do of course each present unique aspects of measurement. Microstructure, assessed via T1w/T2w signal, may offer a shorter acquisition time and thus be preferable for study in certain populations, and is less susceptible to interhemispheric measurement issues. On the other hand, structural connectivity measurements aim to directly assess the strength of existing axonal connections, which microstructural similarity does not do. In this work, we chose to study microstructural similarity in order to test the hypothesis of inter-individual variability in the relationship between function and cyto- and myeloarchitectonic features. Regardless, further efforts to use specific terminology in order to distinguish between these complementary measures should be promoted.

### 4.4 NMF Technical Considerations

NMF is a matrix decomposition technique which has been used in a number of neuroimaging applications to investigate inter-individual differences (Nassar et al., 2018; Patel, Steele, Chen, Patel, Devenyi, Germann, Tardif, & Chakravarty, 2020; Sotiras et al., 2015). Applied here to microstructure-function coupling data, we use NMF to decompose group level coupling measurements into separate additive components, each of which represent a set of brain regions with shared microstructure-coupling variation and are associated with a set of subject specific weightings used to assess inter-individual differences within a given set of brain regions.

A key design decision is that, in contrast to previous neuroimaging applications (Nassar et al., 2018; Patel, Steele, Chen, Patel, Devenyi, Germann, Tardif, & Chakravarty, 2020; Sotiras et al., 2015, 2017; Varikuti et al., 2018), we chose not to enforce orthogonality in the output spatial components. Orthogonality constraints on *W* ensure that there is negligible spatial overlap across components, which can benefit clustering applications (Patel, Steele, Chen, Patel, Devenyi, Germann, Tardif, & Chakravarty, 2020), and aid in interpretation as each brain region participates in only one output component/pattern. Conversely, in this work we apply no such constraint under the assumption that the microstructural and functional variation of interest is more smoothly varying than that conveyed by a hard cluster boundary. Both histological and *in vivo* findings support this notion across multiple data types (García-Cabezas et al., 2019, 2020; Huntenburg et al., 2018). Further, we also aim to uncover multiple patterns of inter-individual variability occurring in the same regions, which is difficult to do if components must be orthogonal. The implication of this is evident when considering the first and third components identified by our analysis. These occupy similar brain regions (**Figure 3**), but analysis of subject weightings show that different subsets of individuals can be identified based on coupling patterns in the same regions. A potential downside of this approach is added complexity, as it is more difficult to definitively describe the features of a given region in comparison to an orthogonal or clustering approach. Thus, while each specific study should consider its own needs and goals, here the loosening of spatial orthogonality constraints was better suited to the specific goal of identifying inter-individual variability in microstructure-function coupling.

### 4.5 Limitations

The primary limitations of this study are related to the correlative nature of the measurements and methods involved. The MRI signals used in this study are sensitive to a wide range of biological phenomena (Tardif et al., 2016; Zatorre et al., 2012). This sensitivity increases the range of phenomena one may query, but the lack of specificity makes it difficult to draw any conclusive statement of cellular level phenomena driving our results. The discussion in above on T1w/T2w as a microstructure measure, as opposed to myelin, is an example of this. Furthermore, despite the high resolution data and equivolume surface methods used, measurements of T1w/T2w sampled at multiple cortical depths may be susceptible to partial volume effects. To study functional connectivity we use a Pearson’s correlation. This is a standard and effective approach to measure the temporal relationship between two regions (Power et al., 2014), though more rigorous metrics such as partial correlation are less susceptible to indirect connections than the method employed here. Our analysis does not incorporate the impact of distance between regions on MPC, or RSFC measurements. While the distance between two regions is heavily related to both measures, previous works have shown that the major dissociations between transmodal and primary cortices hold even when accounting for these measures (Huntenburg et al., 2017). Nonetheless, distance may still be impacting the relationship between MPC and RSFC. We summarized microstructural-functional relations at the node level, as opposed to analyzing each pair of regions. This decision makes the current analysis significantly less complex in terms of both computation and interpretation, and is in line with the approach throughout the field (Huntenburg et al., 2017; Suárez et al., 2020; Valk et al., 2022). However, the downside is we are unable to identify specific sets of connections, or specific pairs of regions, which drive variability. To study the relationship between coupling variability and cognitive performance, we employ PLS. While PLS is able to identify multivariate patterns, it lacks directionality in its analysis. Thus, we are unable to make any inferences on which of microstructure-function coupling variability or cognitive performance is the driving factor in the identified analysis.

### 4.7 Conclusion

We studied inter-individual variability in microstructural-functional coupling in a sample of healthy young adults. Our analysis identified four components, each of which identified inter-individual variability in a select set of cortical regions. We applied a cytoarchitectonic atlas to show that the components distinguished between known cortical types. Further, analysis of subject-specific loading coefficients enabled identification of subsets of individuals with varying coupling patterns, and a latent factor relating coupling strength in idiotypic, limbic, and heteromodal regions to cognitive performance across a number of domains. Our results lend support to histological and *in vivo* findings of regionally varying microstructure-function patterns, and also suggest inter-individual variability in this key brain organization principle is worthy of future study in relation to cognition and disease.

## Acknowledgements

This work was supported by funding from the Fonds de Recherche du Québec Santé, CIHR, NSERC and McGill University (Shuk-Tak Liang Fellowship). RP receives salary support from the Fonds de Recherche du Québec Santé. Dr. Chakravarty receives salary support from the Fonds de Recherche du Québec Santé. Data were provided by the Human Connectome Project, WU-Minn Consortium (Principal Investigators: David Van Essen and Kamil Ugurbil; 1U54MH091657). Data processing was performed in part using the General Purpose Cluster resources of the SciNet HPC Consortium (Loken et al., 2010).

