## Supplementary Materials for "Investigating individual variability in microstructural-functional coupling in the human cortex"

### 1.0 Methods

HCP MRI data was collected on a Siemens 3T Skyra scanner with a custom 100 mT/m gradient (Van Essen et al., 2013, 2012). We used the T1w and T2w images (0.7mm isotropic voxels) and the subsequently derived T1/T2 maps from the HCP project that we used to index myelin concentration. fMRI data was acquired at 3T using an optimized multiband acquisition which greatly reduces acquisition time and enables a resolution of 2 mm isotropic voxel size and high temporal resolution (Glasser et al., 2013; Smith et al., 2013; Uğurbil et al., 2013; Van Essen et al., 2012) (see Supplementary materials, Section 1.1 for acquisition details).

#### 1.1 Magnetic Resonance Imaging data acquisition parameters

T1- and T2-weighted magnetic resonance imaging (MRI) data were used from the HCP Young Adult dataset (David C. Van Essen et al., 2013; D. C. Van Essen et al., 2012). These data were acquired with the following sequence parameters : TR = 2400 ms, TE = 2.14 ms, TI = 1000 ms, flip angle = 8 degrees, bandwidth = 210 Hz/pixel, echo spacing = 7.6 ms, FOV = 224mm, matrix =320✕256. Two separate images were acquired and fat suppression was achieved using a non-selective binomial water excitation pulse. T2w images were similarly acquired at 0.7 mm isotropic resolution using a Siemens SPACE variable flip angle turbo spin-echo sequence, and with sequence parameters as follows: TR = 3200 ms, TE = 565 ms, bandwidth = 744 Hz/pixel, FOV = 224mm, matrix =320✕256. The two separate images were acquired and averaged (Glasser et al., 2013; D. C. Van Essen et al., 2012). Resting state functional MRI (rsfMRI) data were acquired with the following parameters: multiband factor = 8, TE = 33ms, TR = 0.72s, FOV = 208 mm × 180 mm, matrix = 104 × 90. During resting state acquisitions, participants are instructed to keep their eyes open and fixate on a cross hair. Two separate 15 minute sessions with opposing phase encode directions are used in this study (Glasser et al., 2013; Smith et al., 2013; Uğurbil et al., 2013; D. C. Van Essen et al., 2012).

#### 1.2 Generation of T1w/T2w myelin maps

T1w and T2w images underwent preprocessing conducted by the HCP and included gradient distortion correction and co-registration of available repeated runs and averaging. An AC-PC alignment to MNI space was conducted, followed by brain extraction and correction for readout distortions. After applying these corrections, the corrected T2w image was co-registered to the T1w image to create the T1w/T2w ratio image and bias field corrected (Glasser et al., 2013; Glasser & Van Essen, 2011).

### 1.3 Nonegative Matrix Factorization

In order to decompose any *m x n* input matrix (m = rows, n = columns), NMF outputs a *m x k* component matrix, *W*, and a *k x n* weight matrix, *H*. Here *k* represents the number of components, or granularity, of the decomposition and is a user defined parameter. As the name suggests, NMF requires both input and output data to be non-negative. The component and weight matrices are constructed such that their product reconstructs the given input data as best as possible. Thus, NMF represents the input data as a set of basis vectors (components) which recapitulate the input data through a weighted linear combination (weights). NMF seeks to minimize the following objective function:

$$||X - WH||^2 \tag{S1}$$

Where $||\ ||^2$ represents the Frobenius norm. *H* contains subject weightings which can be used to describe the relevance of a given component in the reconstruction of a given subject's coupling measurements. NMF seeks to construct *W* and *H* such that their multiplication represents the original input as best as possible.

### 1.4 Partial Least Squares Analysis

PLS analysis has been described in detail elsewhere (Krishnan et al., 2011; Zeighami et al., 2017). Briefly, PLS analysis begins by computing the correlation matrix between brain and behaviour variables. The correlation matrix is then subject to singular value decomposition which gives three outputs - a matrix of left singular vectors, a matrix of right singular vectors, and singular values. Each pair of singular vectors weight the respective brain and behaviour variables such that they maximally covary, and the associated singular value quantifies the proportion of covariance explained between the brain and behaviour variables. Thus, each LV contains a triplet of information - singular vectors describing the contribution of each brain variable, singular vectors describing the contribution of each behaviour variable, and a singular value denoting the effect size of the LV (Krishnan et al., 2011; McIntosh and Lobaugh, 2004; Zeighami et al., 2017). Statistical significance of LVs is assessed via permutation testing, whereby the input brain data is permuted 5000 times and PLS performed with each permuted dataset in order to create a null distribution of singular values. From this null distribution, a non parametric p value can be obtained for each LV. We used a threshold of $p<0.05$ to determine significance. Bootstrap resampling is used to test the reliability of contribution of brain and behaviour variables. 5000 bootstrapped datasets are created via random sampling with replacement of both brain and behaviour data. PLS is performed on each bootstrapped dataset to obtain distributions of the singular vector weights for each brain and behaviour variable. For brain variables, a bootstrap ratio (BSR) is computed by dividing the singular vector weight by its standard error obtained via the bootstrapped distribution. Given a normal distribution, this is analogous to a z-score, such that a BSR of 2.58 corresponds to a p value of 0.01. For behaviour variables, the bootstrapped distribution is used to obtain a 95% confidence interval for each singular vector weighting associated with the original PLS run on the full dataset (Krishnan et al., 2011; McIntosh and Lobaugh, 2004; McIntosh and Mišić, 2013; Zeighami et al., 2017).

### 1.5 Transcriptomic Analyses: Mean-Centred PLS (mcPLS)

A key difference between mcPLS and the previously applied behavioural PLS is that in mcPLS, data in $X$ and $Y$ are arranged by group. This makes mcPLS a preferred technique for identifying relationships between groupings in a set of variables. Here, $X$ is created by vertically concatenating each component samples x genes matrix. $X$ is thus a whole brain expression matrix, with each component a group and each sample an observation. $Y$ is a binary matrix encoding component membership for each sample. We then compute a matrix $M$, by taking the component-wise average within each gene (i.e. column) of $X$. $M$ is then mean-centered in a column-wise fashion and submitted to a singular value decomposition to output LVs which describe gene expression patterns that separate the input components (i.e. groups). LVs contain a singular value describing the proportion of covariance explained, gene weights which describe the expression pattern of the LV, and contrast values which describe the degree to which a given component is associated with the LV expression pattern.

**Table S1** lists the tests from the HCP behavioural testing which were included in the present study. The left column denotes the abbreviation/description used in the main text, the middle column denotes the instrument used for testing, and the right column denotes to cognitive subdomain, as described in Barch et al., 2013 and at https://wiki.humanconnectome.org/display/PublicData/HCP-YA+Data+Dictionary-+Updated+for+the+1200+Subject+Release. Abbreviations: TC: Total correct responses; SI: Skipped items; RTCR: Median reaction time for correct responses; CRTE: Median reaction time divided by expected number of clicks for correct trials; OFF: Total positions off for all trials.

| Abbreviation used in Main Text | Test/Instrument | Cognitive Subdomain |
|---|---|---|
| Episodic Memory | NIH Toolbox Picture Sequence Memory Test: Age-Adjusted Scale Score | Episodic Memory |
| Cognitive Flexibility | NIH Toolbox Dimensional Change Card Sort Test: Age-Adjusted Scale Score | Executive function/cognitive flexibility |
| Inhibition | NIH Toolbox Flanker Inhibitory Control and Attention Test: Age-Adjusted Scale Score | Executive Function/Inhibition |
| Fluid Intelligence CR | Penn Progressive Matrices: Number of Correct Responses | Fluid Intelligence |
| Fluid Intelligence SI | Penn Progressive Matrices: Total Skipped Items | Fluid Intelligence |
| Fluid Intelligence RTCR | Penn Progressive Matrices: Median Reaction Time for Correct Responses | Fluid Intelligence |
| Language Decoding | NIH Toolbox Oral Reading Recognition Test: Age-Adjusted Scale Score | Language/Reading Decoding |
| Language Comprehension | NIH Toolbox Picture Vocabulary Test: Age-Adjusted Scale Score | Language/Vocabulary Comprehension |
| Processing Speed | NIH Toolbox Pattern Comparison Processing Speed Test: Age-Adjusted Scale Score | Processing speed |
| DDisc AUC_200 | Delay Discounting: Area Under the Curve for Discounting of $200 | Self-regulation/impulsivity |
| DDisc AUC_40k | Delay Discounting: Area Under the Curve for Discounting of $40 000 | Self-regulation/impulsivity |
| Spatial Orientation TC | Variable Short Penn Line Orientation: Total Number Correct | Spatial orientation |
| Spatial Orientation CRTE | Variable Short Penn Line Orientation: Median Reaction Time Divided by Expected Number of Clicks for Correct | Spatial orientation |
| Spatial Orientation OFF | Variable Short Penn Line Orientation: Total Positions Off for All Trials | Spatial orientation |

| | | |
|---|---|---|
| Sustained Attention SEN | Short Penn Continuous Performance Test: Sensitivity | Sustained Attention |
| Sustained Attention SPEC | Short Penn Continuous Performance Test: Specificity | Sustained Attention |
| Verbal Episodic Memory TC | Penn Word Memory Test: Total Number of Correct Responses | Verbal episodic memory |
| Verbal Episodic Memory RTCR | Penn Word Memory Test: Median Reaction Time for Correct Responses | Verbal episodic memory |
| Working Memory | NIH Toolbox List Sorting Working Memory Test: Age-Adjusted Scale Score | Working Memory |

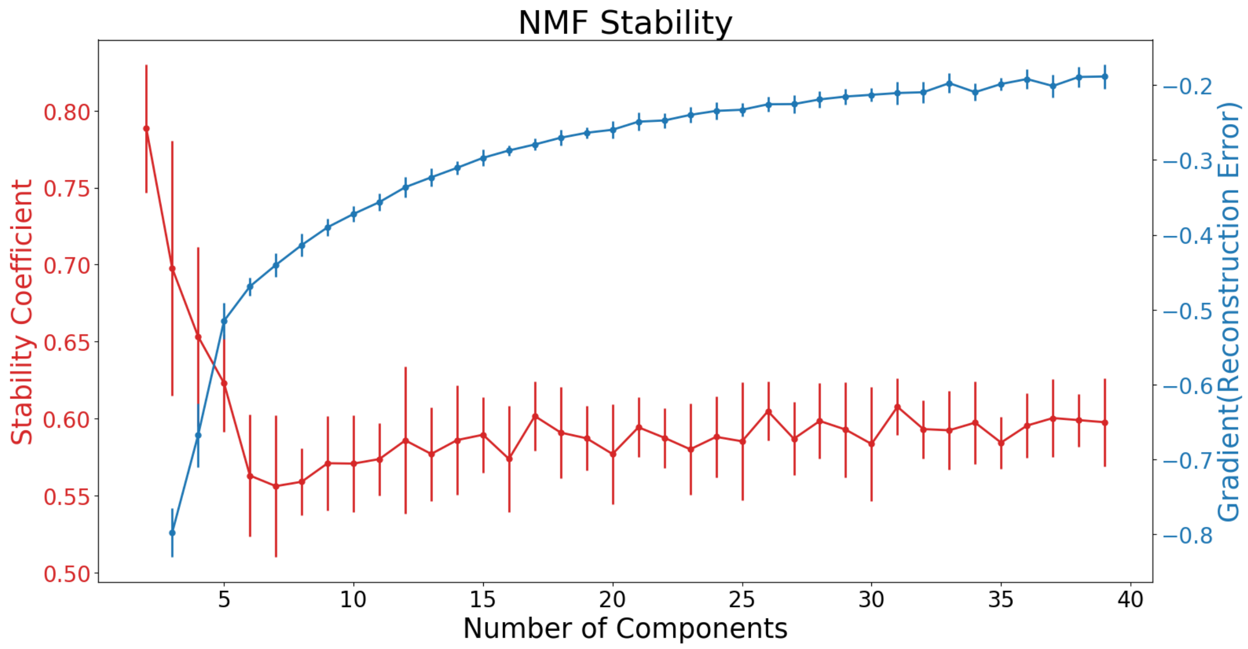


**Figure S1: Stability analysis suggests k=4 for further analysis.** Both the stability coefficient (red) and gradient of the reconstruction error (blue) are plotted on y axis vs number of components on the x axis. A high stability coefficient indicates that the patterns identified are replicable across varying subsets of individuals, while a low coefficient suggests the identified patterns are less consistent across subjects. Large changes in the gradient of reconstruction error indicate that the accuracy of the NMF solution at a given granularity is very different compared to the previous granularity, with smaller changes indicative of marginal gains in accuracy.
